# Modeling the longitudinal changes of ancestry diversity in the Million Veteran Program

**DOI:** 10.1101/2022.01.24.477583

**Authors:** Frank R Wendt, Gita A Pathak, Jacqueline Vahey, Xuejun Qin, Dora Koller, Brenda Cabrera-Mendoza, Angela Haeny, Kelly M Harrington, Nallakkandi Rajeevan, Linh M Duong, Daniel F Levey, Flavio De Angelis, Antonella De Lillo, Tim B Bigdeli, Saiju Pyarajan, VA Million Veteran Program, J. Michael Gaziano, Joel Gelernter, Mihaela Aslan, Dawn Provenzale, Drew A. Helmer, Elizabeth R. Hauser, Renato Polimanti, Department of Veteran Affairs Cooperative Study Program (#2006)

## Abstract

The Million Veteran Program (MVP) participants represent 100 years of US history, including significant social and demographic change over time. Our study assessed two aspects of the MVP: (i) longitudinal changes in population diversity and (ii) how these changes can be accounted for in genome-wide association studies (GWAS). The MVP was divided into five birth cohorts (N-range=123,888 [born from 1943-1947] to 136,699 [born from 1948-1953]). Groups of participants were defined by (i) HARE (harmonized ancestry and race/ethnicity) and (ii) a random-forest clustering approach using the 1000 Genomes Project and the Human Genome Diversity Project (1kGP+HGDP) reference panels (77 world populations representing six continental groups). In these groups, we performed GWASs of height, a trait potentially affected by population stratification. Birth cohorts demonstrate important trends in ancestry diversity over time. More recent HARE-assigned Europeans, Africans, and Hispanics had lower European ancestry proportions than older birth cohorts (0.010<Cohen’s d<0.259, p<7.80×10^−4^). Conversely, HARE-assigned East Asians showed an increase in European ancestry proportion over time. In GWAS of height using HARE assignments, genomic inflation due to population stratification was prevalent across all birth cohorts (linkage disequilibrium score regression intercept=1.08±0.042). The 1kGP+HGDP-based ancestry assignment significantly reduced the population stratification (mean intercept reduction=0.045±0.007, p<0.05) confounding in the GWAS statistics. This study provides a comprehensive characterization of ancestry diversity of the MVP cohort over time and highlights that more refined modeling of genetic diversity (e.g., the 1kGP+HGDP-based ancestry assignment) can more accurately capture the polygenic architecture of traits and diseases that could be affected by population stratification.

## Introduction

Genome-wide association studies (GWAS) have successfully identified loci associated with thousands of human traits and diseases using extremely large sample sizes.^1^ Multi-ancestry cohorts, such as the Department of Veterans Affairs (VA) Million Veteran Program (MVP), offer unique opportunities to study the genetic architecture of complex traits across diverse populations.^2^ As of June 2021 MVP has enrolled more than 840,000 Veteran volunteers, >650,000 of whom have been genotyped, and includes a wide range of phenotypic and health outcome information. Generally, GWAS are conducted within samples stratified by genetically-determined ancestry groups and using genetic principal components to account for within-ancestry population structure.^3^ Recently, other methods have been proposed to improve the modeling of genetic diversity and the gene discovery of complex traits in diverse populations.^4–6^ With respect to MVP diversity classification, the HARE (harmonized ancestry and race/ethnicity) approach was developed to inform genetic ancestry assignments by leveraging self-identified racial and ethnic (SIRE) background under the hypothesis that these variables provide complementary information and may improve the appropriateness of population strata in genetic studies.^3^ The HARE approach uses supervised machine learning and genetically determined ancestry to refine SIRE information for GWAS in three ways: (i) identify individuals whose SIRE is inconsistent with genetic information, (ii) reconcile conflicts among multiple SIRE sources, and (iii) impute missing racial/ethnic information when the predictive confidence is high. Although the HARE approach aims to increase the inclusivity of the population group definition to reduce the number of unclassified individuals, it can also, by the same process, increase the heterogeneity and the complexity of genetic structure within each HARE-defined group. Additionally, the inclusion of SIRE information can introduce biases related to specific racial and ethnic classifications used. For example, SIRE information used in MVP is based on racial and ethnic categories defined by the US Census. In this classification, the “Asian” group includes two distinct ancestry groups – Central/South Asian and East Asian – that are very different from a genetic perspective.^7^ Therefore, a GWAS conducted on a HARE-assigned “Asian” superpopulation has a high risk to be biased by population stratification unaccounted for by genetic principal components. To a lesser extent, population stratification could also affect GWAS conducted in samples defined using HARE assignment due to an increased genetic heterogeneity.

To test this hypothesis, we compared the HARE approach with a classification based on genetic ancestry categories derived from a high-resolution reference panel (77 world populations representing six continental groups),^8; 9^ testing how they model genetic diversity in a GWAS of height. Previous studies demonstrated that height polygenic architecture can be strongly affected by unaccounted for population stratification.^10^ To understand multiple scenarios related to the different sample compositions over time that characterize the US and therefore the MVP, we stratified the MVP cohort, which spans almost 100 years (from 1904-1999), into five birth cohorts of approximately 130,000 MVP participants each. Consistent with US demographics and changes in military policies, the demographic characteristics of US military personnel have changed drastically over time with more personnel identifying as Black, Hispanic, Asian, or other non-European descent categories in more recent decades.^10; 11^ Accordingly, the five birth cohorts will present different ancestry compositions reflecting these social and demographic changes. But they also reflect demographic changes reflected in differing admixture in ancestry groups over time and social changes in self-identification. These cohorts permitted us to assess how different superpopulation-assignment approaches work in different scenarios to correct the population stratification affecting height polygenic architecture.^10^ This trait was selected due to the well-documented unaccounted-for effects of population stratification in large genetic studies.^10^ Our findings provide a comprehensive evaluation of the challenges in accurately modeling the diversity of human populations in the context of multi-ancestry GWAS. Additionally, we characterized the longitudinal changes of ancestry composition in the MVP cohort, showing how social and demographic changes can affect the genetic structure and introduce specific challenges in the design of GWAS. This study presents one possible solution, a higher resolution ancestry reference panel, to mitigating these effects on genetic structure in GWAS.

### Subjects and Methods

#### Definitions of Race, Ethnicity, and Ancestry Groups

This study compares different cohorts of Veterans grouped together based on genetic data or on HARE. HARE relies on the blending of self-identity and genetic information; we will use the following terminology for clarity. “Superpopulation” to reference a group of participants defined by HARE or genetic data. “Ancestry” is strictly applied to population defined by genetic data only. “Ethnicity” describes a population of people with common national and/or cultural traditions. In the MVP, participants self-reported one of the following ethnicities: “not Spanish, Hispanic, or Latino,” “Mexican, Mexican American, or Chicano,” “Puerto Rican,” “Cuban,” or “Other Spanish, Hispanic, or Latino.”^2^ Finally, “race” is a social construct that groups individuals by self-identity and encompasses many aspects of cultural belonging and physical appearance. In the MVP, participants self-reported one or more of the following races: “White,” “Black or African American,” “Chinese,” “Japanese,” “Asian Indian,” “Other Asian,” “Filipino,” “Pacific Islander,” and/or “Other.”^2^ Populations grouped by ancestry, race, or ethnicity generally overlap in the MVP (e.g., non-Spanish, Hispanic, or Latino Black or African American Veterans generally have high proportions of continental African ancestry).

#### Cohort Description

The MVP is an ongoing voluntary research cohort of the United States military population composed of active users of the Veterans Health Administration healthcare system who learn of the MVP by invitational mailing and/or from MVP staff while receiving clinical care. All MVP participants provided informed consent and Health Insurance Portability and Accountability Act (HIPAA) of 1996 authorization. As of 2021, approximately 840,000 Veterans have enrolled in the program.^12^ Research involving the MVP data was approved by the Veterans Affairs (VA) Central Institutional Review Board (IRB). The current project was also approved by VA IRBs in Durham (North Carolina), Houston (Texas), Boston (Massachusetts), and West Haven (Connecticut).

The MVP integrates data from the Electronic Health Record and at least two surveys administered at the time of recruitment.^2^ The Baseline Survey collects data regarding demographics, family pedigree, health status, lifestyle habits, military experiences, medical history, family history of illness, and physical features. The Lifestyle Survey asks questions from validated instruments in domains selected to provide information about sleep and exercise habits, environmental exposures, diet, and sense of well-being.

#### SNP Genotyping and Quality Control

For the current analysis, we used the release 4 data freeze consisting of genotype data available for 658,582 participants (8.9% females and 29.4% from non-EUR HARE superpopulations). Genotyping was performed with the MVP 1.0 custom Axiom® Biobank array consisting of 668,418 SNP assays. The details on the quality control, superpopulation assignment, relatedness, and imputation have been described previously.^12^

#### Harmonized Ancestry and Race/Ethnicity (HARE)

HARE was designed to define strata for superpopulation-specific GWAS using a two-stage categorization procedure.^3^ First, a support vector machine (SVM) was built to learn the correspondence between genetically determined ancestry and SIRE information. Second, HARE was assigned based on the harmonization of SIRE, genetically determined ancestry, and the trained SVM. There are four HARE superpopulations in MVP Release 4: non-Hispanic Black with predominantly African ancestry (N=123,120), non-Hispanic Asian with predominantly East Asian ancestry (N=8,329), non-Hispanic white with predominantly European ancestry (N=464,961), and Hispanic with predominantly Admixed American ancestry (N=52,183). HARE unclassified status can be due to (i) lack of SIRE and predicted probability of ancestry cannot be resolved between two populations or (ii) discordance between genetic information and SIRE.^3^ A total of 9,989 participants (1.52%) could not be classified by HARE.

#### High-resolution Ancestry Reference Panel

A high-resolution ancestry reference panel of non-MVP individuals was created following procedures developed in the Pan-ancestry UK Biobank initiative (see https://pan.ukbb.broadinstitute.org/ for full details). The reference panel combines individuals from the 1000 Genomes Project (1kGP Phase 3; 26 populations across five continental ancestries)^13^ and Human Genome Diversity Project (HGDP; 51 populations across six continental ancestries).^9^ Hereafter this panel is referred to as 1kGP+HGDP. These data were stratified into continental ancestries according to their recruitment strategies and the previous genetic inference analyses.^9; 13^ The ancestry groups include African (AFR), Central/South Asian (CSA), East Asian (EAS), European (EUR), Middle Eastern (MID), and Admixed American (AMR). SNP coordinates were assigned relative to the hg37 reference genome. After quality control for minor allele frequency (1%), missingness (SNP=5% and individual=3%), heterozygosity, and pairwise kinship estimates, the high-resolution ancestry panel included 3,284 individuals. Principal components analysis (PCA) was performed on unrelated individuals from the reference panel. First, we applied PCA on a high quality and linkage disequilibrium (LD)-pruned (R^2^=0.01, 1500-kb window size) dataset of genotypes from the MVP using plink 2.0.^14^ The PC loadings from top 20 PCs of the 1kGP+HGDP reference panel (training data) were projected onto PC loadings of genotypes from the MVP participants. Ancestry assignments were performed using the random forest classifier implemented in the `randomForest` R package with default settings. The projected ancestry groupings in MVP were further refined for ancestry outliers using the top 20 PCs within the assigned ancestry group for the MVP cohort. We defined outliers based on PC loadings outside six median absolute deviations across the first 20 PCs.

#### Birth Cohort Definition

Each MVP participant endorsed service in one of nine possible service eras: 1941 or earlier, December 1941 to December 1946, January 1947 to June 1950, July 1950-January 1955, February 1955 to July 1964, August 1964 to April 1975, May 1975 to July 1990, August 1990 to August 2001, or September 2001 or later.^2^ Though MVP participants are mapped to service era, these strata represent overlapping periods of service with >22% of MVP participants serving in multiple eras (Table S1), 4.3% of whom served in non-contiguous eras. For these reasons, military service data were not used to stratify the MVP. Using self-reported participant birth year from the MVP Baseline Survey,^2^ we used a cumulative distribution function to identify approximately equally sized birth cohorts (BCs). Each BC consisted of approximately 130,000 participants (Table S2).

#### Height as a Model Trait to Investigate Population Stratification

This study aims to detect and quantify the effect of residual population stratification among ancestry assignments. GWASs of height show severe biases attributed to unaccounted-for ancestry diversity among discovery samples.^10^ Height in the MVP was measured in inches as part of the core vital signs assessment at participant enrollment. GWAS of height were performed using age, sex, and 10 within-superpopulation PCs as covariates.^10; 12; 15^ Covariate PCs were calculated per superpopulation per classification method resulting in ancestry- and method-specific PCs for each HARE superpopulation and each 1kGP+HGDP ancestry group.

#### Ancestry Proportion

The ancestry proportions among MVP participants were estimated using ADMIXTURE.^16^ With ADMIXTURE, each MVP participant was assigned five ancestry proportions using reference populations from 1kGP Phase 3: Han Chinese in Beijing (CHB), British in England and Scotland (GBR), Luhya in Webuye, Kenya (LWK), Peruvian in Lima, Peru (PEL), and Yoruba in Ibadan, Nigeria (YRI). These reference populations were selected to represent homogeneous continental ancestries along with a relatively large Admixed American reference population (PEL).

#### Detection of Unaccounted-For Ancestry Diversity

The outcome used to quantify the presence of unaccounted-for ancestry diversity in each GWAS was the linkage disequilibrium score regression (LDSC) intercept and attenuation ratio.^17^ The LDSC intercept assesses whether the distribution of genome-wide association statistics is consistent with an expected distribution. The LDSC intercept of GWAS typically range from 1-1.05 with values greater than 1.05 often considered as evidence of systematic bias in the test statistics.^17–19^. Attenuation ratios quantify the proportion of inflation in the mean χ^2^ statistic that can be ascribed to causes other than polygenicity. When the LDSC intercept is in an acceptable range, the attenuation ratio typically ranges from 0-20%. As the LDSC intercept exceeds 1.05, attenuation ratios in this range indicates the presence of confounding, typically attributed to unaccounted-for ancestry diversity. Two-sided Z-tests were used to compare the statistical difference in LDSC intercepts and attenuation ratios between groups. Multiple testing correction was applied to these results using the false discovery rate (FDR < 5%).

## Results

### Birth Cohort Description

Birth years among MVP participants span 96 years (1904-1999). Five birth cohorts were defined, each consisting of approximately 130,000 participants: BC1 1904-1942 (N=131,587), BC2 1943-1947 (N=123,888), BC3 1948-1953 (N=136,699), BC4 1954-1963 (N=127,414), BC5 1964-1999 (N=133,513). The distribution of HARE superpopulations and service era descriptive statistics are shown in Tables S1 and S2. In line with US military population demographic shifts, more contemporary birth cohorts included more females and members of non-European ancestry.

### Longitudinal Changes in Ancestry Diversity

Using PCs calculated within HARE superpopulations, observational changes are seen in the two-dimensional projections of participants’ genetic diversity. Consistent with conventional expectations of two-dimensional PC space, clustering of the first two HARE PCs separates EUR, EAS, and AFR continental ancestry groups. When projected as independent birth cohorts (Figure 1), we show that over time, areas of two-dimensional feature space occupied by genetically heterogeneous individuals become more populous while continental population clusters become more heterogeneous. In other words, admixture increased over time and the younger cohorts were more genetically heterogenous than the older ones Across different projections of feature space (Figure 1), the most recent cohort (BC5 1964-1999) clearly shows a greater separation of continental groupings and a larger number of Hispanic and unclassified participants.

**Figure 1.**
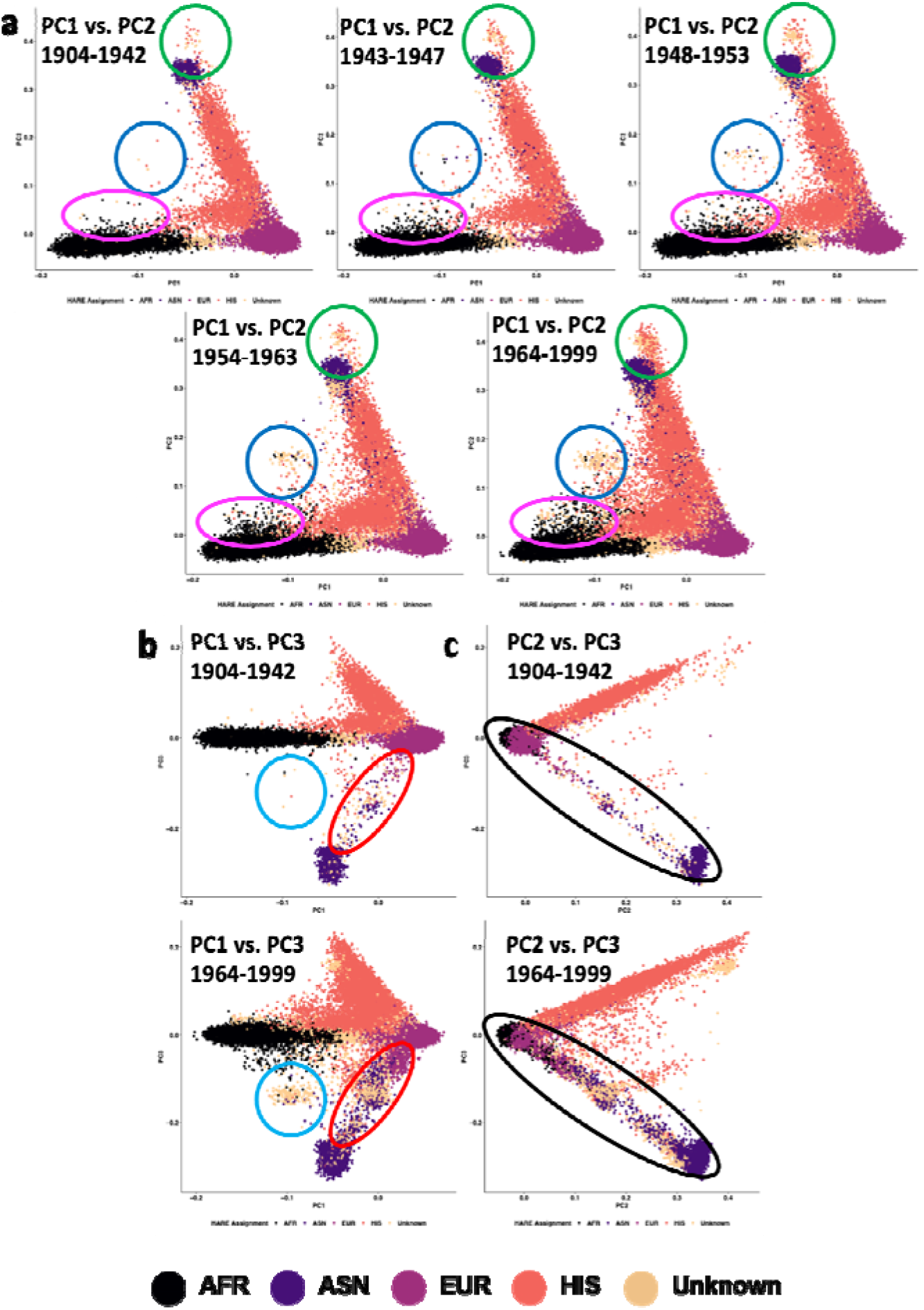
Observational changes in HARE superpopulation heterogeneity across birth cohort in the MVP. Each plot represents an independent birth cohort of approximately 130,000 participants. Panel (a) projects PCs 1 and 2 into two-dimensional space for each birth cohort. In panels (b) and (c), the oldest and youngest birth cohorts are projected on PC1-versus-PC3 and PC2-versus-PC3, respectively, demonstrating comparable observations when using different combinations of PCs. Clusters of participants are circled to draw attention to observational changes in population heterogeneity across birth cohort. The colors differences used to identify these regions are arbitrary but permit easy tracking of highlighted regions across plots. Abbreviations: African (AFR); Asian (ASN); European (EUR); Hispanic (HIS).

We compared ancestry proportions across birth cohorts. Among HARE-EUR participants, there was a decrease in GBR ancestry from oldest to most recent birth cohort while YRI and CHB ancestry proportions increased. While significant due to the large MVP sample size (FDR Q < 0.05), the effect size of these changes among HARE-EUR participants were relatively small (0.01 ≤ |Cohen’s d| ≤ 0.14; Figure 2). HARE-AFR and HARE-HIS groups demonstrated similar significant changes in ancestry proportion across birth cohorts, but these differences also were relatively small. Conversely, HARE-EAS showed a significant decrease in CHB ancestry proportion and an increase in GBR ancestry proportion across birth cohorts. The mean GBR ancestry proportion among HARE-EAS was 3.04% ± 10.2 (BC1 1904-1942), 6.17% ± 14.7 (BC2 1943-1947), 7.76% ± 16.4 (BC3 1948-1953), 8.95% ± 17.0 (BC4 1954-1963), and 9.66% ± 18.1 (BC5 1964-1999). These observations translate to large standardized effect sizes (1.69 ≤ |Cohen’s d| ≤ 3.43). All ancestry proportions and effect sizes are shown in Table S3.

**Figure 2.**
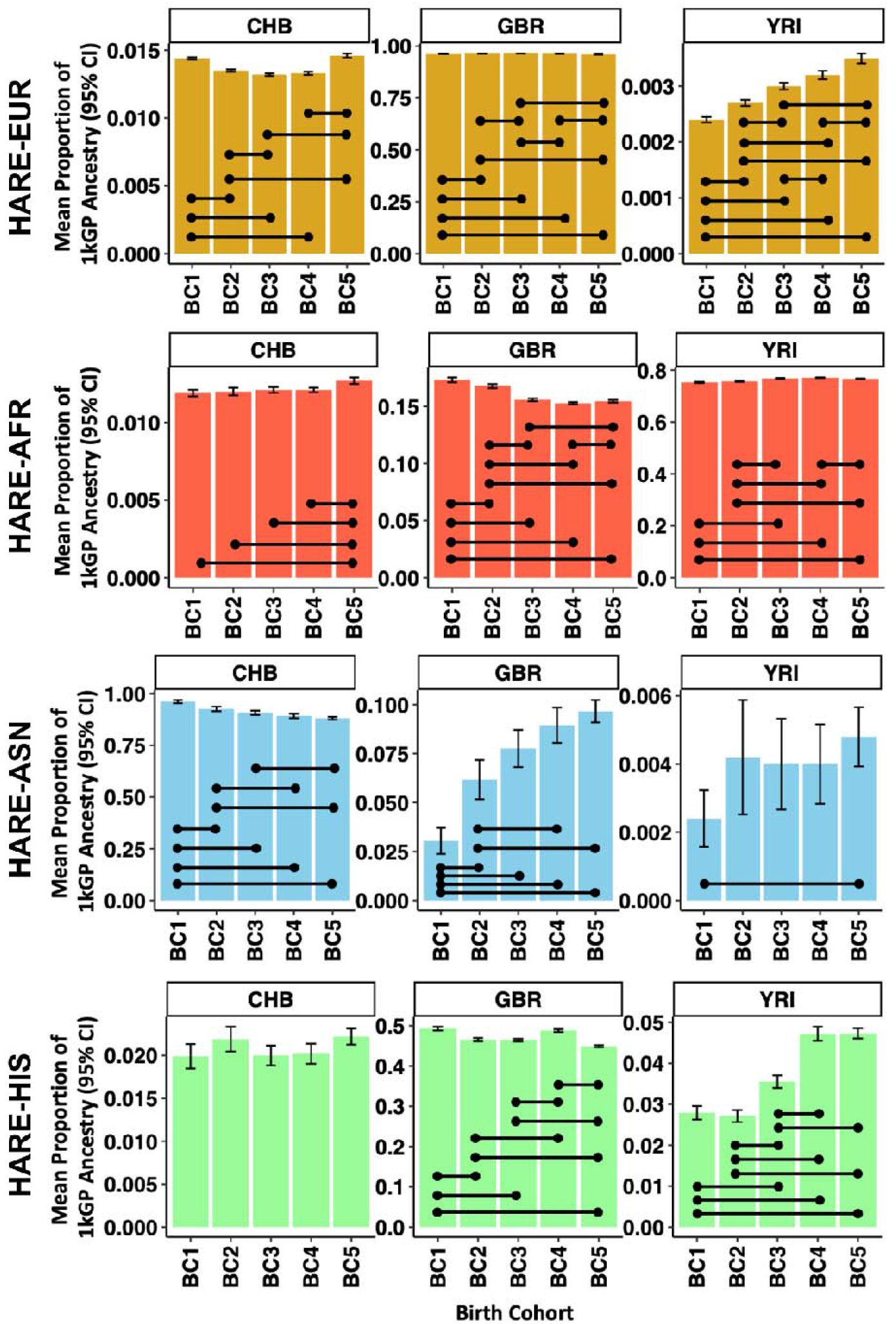
Statistical changes in ancestry diversity in four HARE superpopulations. Each facet per row shows the ancestry proportion for three major continental populations (CHB is an East Asian reference population of Han Chinese in Beijing, China; GBR is a European reference population from Great Britain; YRI is a West African reference population of Yoruba in Ibadan, Nigeria). Birth cohorts are shown across the x-axis and are arranged from oldest (BC1) to most recent (BC5). Black lines connecting colored bars designate significant differences in ancestry proportion between those two birth cohorts (p < 2.0×10^−4^ based on five birth cohorts, five ancestry proportion references, and ten birth cohort pairwise comparisons). All proportions, specific p-values from comparisons, and HARE assignments are shown in Table S3. Abbreviations: African (AFR); Asian (ASN); European (EUR); Hispanic (HIS).

### Height Changes over Time

There were relatively small changes in height across birth cohorts (Table S4). Compared to the oldest birth cohort, more contemporary HARE-AFR individuals were shorter (difference in means = 0.83 inches, Cohen’s *d*=0.23, p=4.26×10^−127^), HARE-ASN individuals were taller (difference in means = 0.96 inches, Cohen’s *d*=0.31, p=4.84×10^−23^), HARE-EUR individuals were taller (difference in means = 0.08 inches, Cohen’s *d*=0.03, p=2.73×10^−7^), and HARE-HIS individuals were taller (difference in means = 0.34 inches, Cohen’s *d*=0.11, p=1.34×10^−14^). In the MVP, the change in GBR ancestry proportion between two birth cohorts, *d*_GBR_ (beta=0.856, p=0.001), was a significant independent correlate of the change in height between the same two birth cohorts (*d*_height_; Figure S1). No significant correlation was observed with respect to other ancestry proportions (p>0.05).

### Distribution of MVP Participants across Ancestry Groups

Using a random forest assignment of ancestry based on the high-resolution reference panel composed of the 1kGP+HGDP (N=3,284 unrelated reference individuals from 77 populations across six continental ancestries), the MVP was stratified into six distinct ancestries (Figure 3). Table 1 shows negligible differences in the numbers of European, African, and Admixed American MVP participants applying the two classification methods. Relative to HARE, the higher diversity ancestry panel applied here permitted the statistical resolution of East Asian, Central/South Asian, and Middle Eastern ancestries. We report a 31.3% increase in the sample size of MVP participants with genetically homogeneous East Asian ancestry. Additionally, the 1kGP+HGDP-based ancestry reduced the number of unclassified individuals compared to the HARE method (4,750 vs. 9,989, respectively).

**Table 1.** Sample size for each superpopulation of the Million Veteran Program cohort using HARE and a high-resolution ancestry panel from the 1000 Genomes Project and Human Genome Diversity Project (1kGP+HGDP).

| Superpopulation group | 1kGP+HGDP-based Random Forest classification | HARE |
| --- | --- | --- |
| European | 459,697 | 464,961 |
| Central/South Asian | 709 | - |
| African | 123,687 | 123,120 |
| Admixed American (referred to as “Hispanic/HIS” by HARE) | 58,034 | 52,183 |
| East Asian (referred to as “Asian/ASN” by HARE) | 10,942 | 8,329 |
| Middle Eastern | 763 | - |
| Unclassified | 4,750 | 9,989 |
| Total | 658,582 | 658,582 |

**Figure 3.**
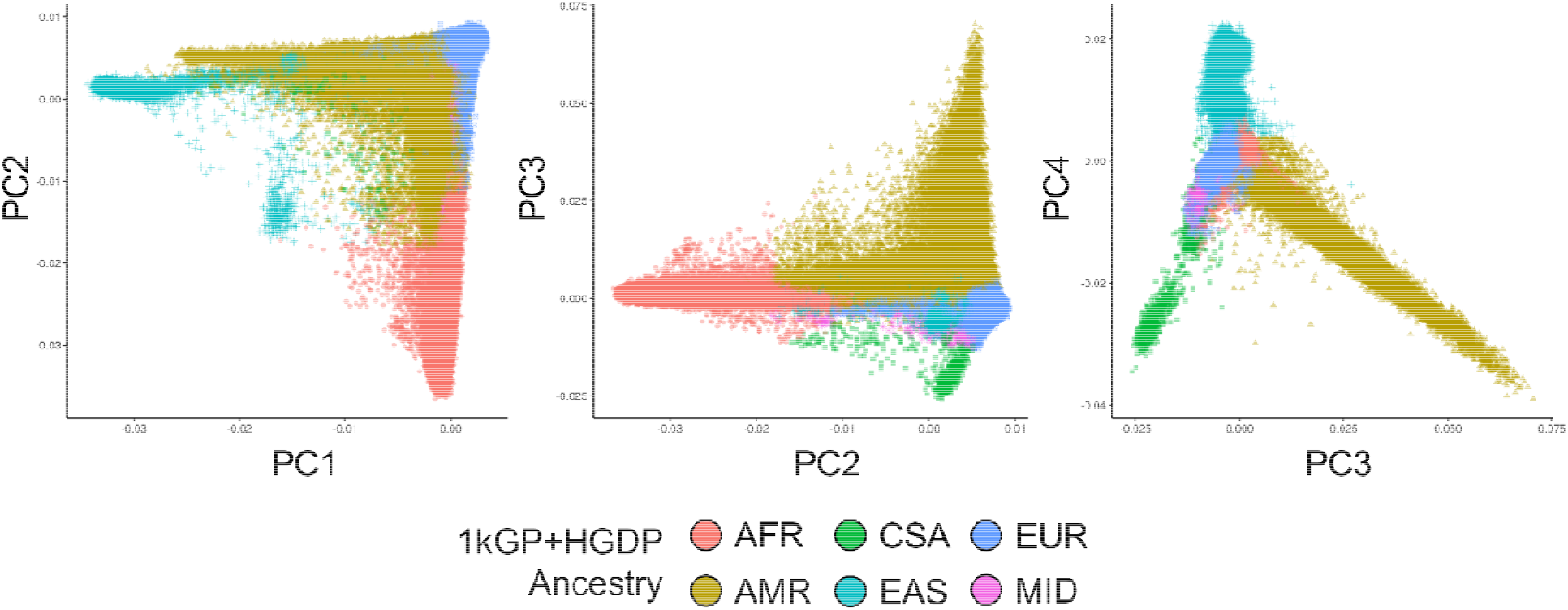
Principal components analysis for genetic ancestry of MVP participants using the random forest classifier method and the 1kGP+HGDP reference panel. Abbreviations: African (AFR), Central/South Asian (CSA), East Asian (EAS), European (EUR), Middle Eastern (MID), and Admixed American (AMR).

### GWAS of Height

We performed GWASs of height in the MVP stratified by BC, ancestry, and ancestry-assignment methods (5 BCs x 4 superpopulations x 2 superpopulation-assignment methods). Due to the lack of a HARE comparator group, height was not assessed in the 1kGP+HGDP Central/South Asian or Middle Eastern ancestries. There were no significant differences in SNP-heritability across methods (Figure S2 and Tables S5 and S6). Among HARE superpopulations, 14 of 20 LDSC intercepts from GWAS of height were greater than 1.05 (HARE-EUR mean intercept=1.13±0.02, HARE-AFR mean=1.07±0.04, HARE-ASN mean=1.06±0.03, and HARE-HIS mean=1.05±0.02; Figure 4). Accounting for population stratification using the high-resolution 1kGP+HGDP ancestry reference panel reduced the LDSC intercept of 18 of 20 GWAS, reflecting lower effects of confounding by population stratification on these GWAS relative to HARE assignments. In four analyses, the LDSC intercept of height GWAS was significantly lower (p < 0.05) in the 1kGP+HGDP population relative to the HARE superpopulation assignment: EUR 1954-1963 (HARE intercept = 1.12 ± 0.017, 1kGP+HGDP intercept = 1.07 ± 0.016, p_diff_ = 0.033), EUR 1964-1999 (HARE intercept = 1.11 ± 0.018, 1kGP+HGDP intercept = 1.06 ± 0.016, p_diff_ = 0.034), EAS 1964-1999 (HARE intercept = 1.10 ± 0.010, 1kGP+HGDP intercept = 1.05 ± 0.010, p_diff_ = 0.003), and HIS 1943-1947 (HARE intercept = 1.07 ± 0.010, 1kGP+HGDP intercept =1.03 ± 0.010, p_diff_ = 0.008). There was no difference in attenuation ratio across methods suggesting that the proportion of test statistic inflation attributable to ancestry stratification has not changed although the high-resolution reference panel accounts for this ancestry diversity significantly better than HARE assignments (Figure S3).

**Figure 4.**
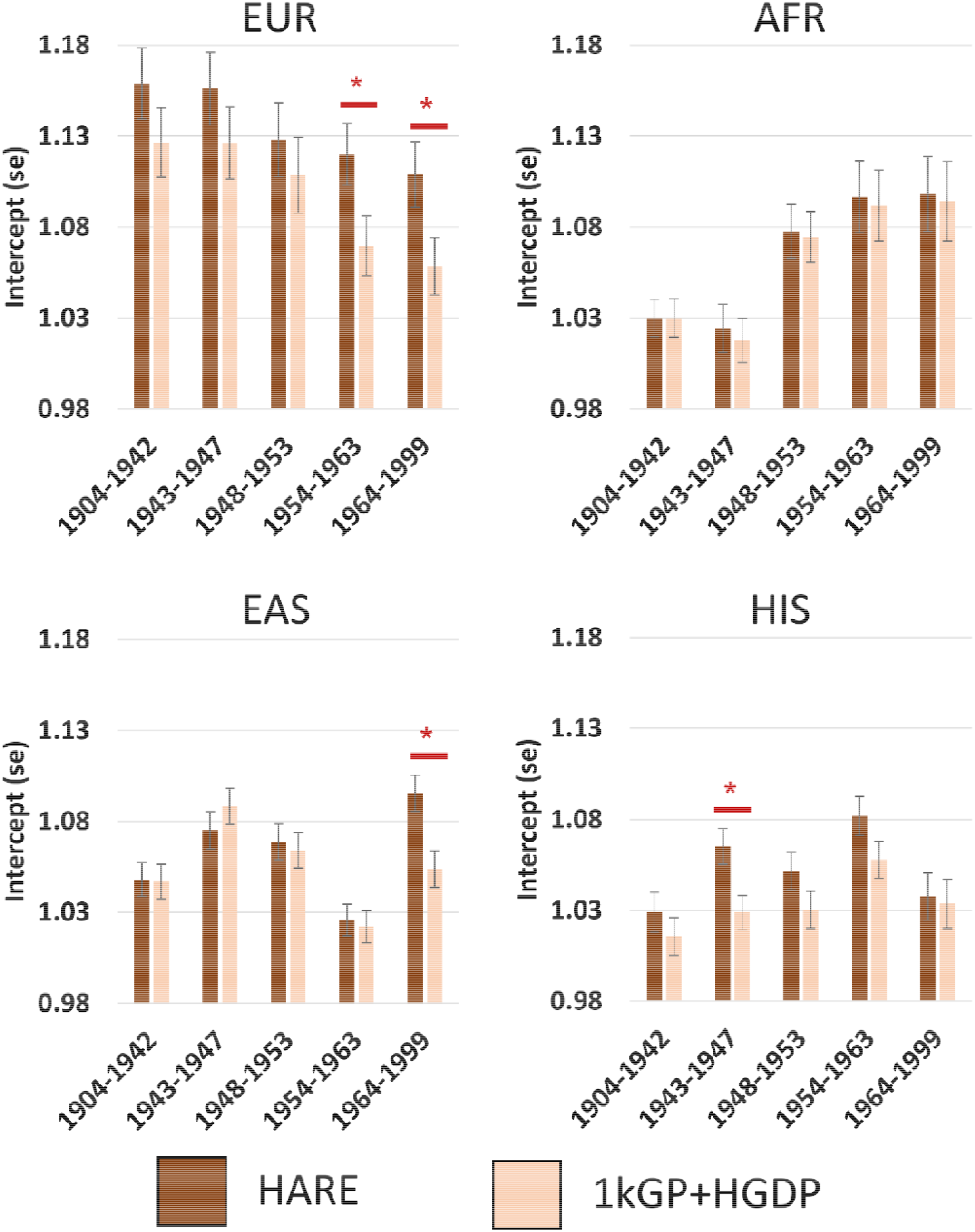
LDSC intercept comparisons across GWAS of height performed in HARE superpopulations and populations assigned using a high-resolution ancestry reference panel composed of 1000 Genomes Project plus Human Genome Diversity Project individuals (1kGP+HGDP). Red asterisks indicate significant difference in intercept estimates (P < 0.05). Each GWAS was performed in unrelated participants of the indicated ancestry with age, sex, and 10 within-population principal components as covariates. Abbreviations: European (EUR); African (AFR); East Asian (EAS); Hispanic (HIS).

## Discussion

People of diverse backgrounds have been historically excluded from genetic studies of health and disease stemming from several social, political, ideological, scientific, and practical factors.^20^ Because of this disproportionate recruitment of study participants, many efforts are now being employed to diversify genetic data collection and make better use of existing data from genetically diverse populations.^21–23^ However, accurate modeling of human genetic variation is essential for unbiased gene discovery in diverse populations.^5; 24–26^ One approach to accomplish this goal this is HARE.^3^ Using machine learning, HARE blends SIRE with genetic data to classify individuals for which these measures of identity and ancestry align. Based on the notion that self-identified race/ethnicity generally correlates with genetically determined ancestry,^3^ an individual’s HARE and SIRE are identical when SIRE is unambiguous; however, SIRE in the MVP is culturally tuned to the demographics of the United States and may not permit generalization of HARE outside genetics research in the United States. Here, we demonstrated that the use of SIRE by the HARE approach captures population dynamics that do not reflect the ancestry of the participants investigated. Most notably, we quantify a longitudinal reduction in East Asian ancestry among HARE-ASN individuals which likely reflects (i) recent admixture in the most contemporary birth cohort that was not present in the oldest birth cohort and/or (ii) the known inclusion of South Asian participants in the HARE-ASN group.^3^

To evaluate HARE, we used a more objective comparator that did not rely on self-report at all: a different approach to cluster MVP participants according to their genetically defined ancestry with a high-resolution reference panel. In GWAS of height based on 1kGP+HGDP ancestry assignment, we identified significant reductions in LDSC intercepts when compared to intercepts obtained from the GWAS conducted using HARE superpopulation assignments. The differences were most pronounced among more recent EUR and EAS populations and may be due to longitudinal changes in ancestry proportion over the five birth cohorts spanning almost 100 years. MVP participants in HARE-EUR and HARE-EAS superpopulations who were born between 1964 and 1999 had a lower proportion of EUR ancestry than earlier birth years. These differences in recent birth cohorts likely indicate that more granular modeling is important to account for recent demographic changes that occurred in the United States. Our results highlight that genetic studies in more genetically diverse cohorts may be more confounded by population stratification when using very broad definitions of ancestry, such as HARE, and that this increased genetic diversity can be modeled better using a high-resolution ancestry reference panel.

A major pitfall to including SIRE categories is that they are based on historical population classifications that are specific to the country(s) where the recruitment and the assessment are performed. MVP SIRE information is based on the classification used by the US Census. This can be very different from those used in other countries and do not reflect the continuum of genetic diversity across human populations. The strongest example is the fact that MVP HARE classification groups together all Asian populations. In the MVP, these include individuals who identify as “Chinese,” “Japanese,” “Asian Indian,” “Other Asian,” or “Filipino,” but Asian populations are extremely heterogeneous, with the largest genetic differences between Central/South Asia and East Asia. Another important difference is the classification of ethnicity, which is specifically related to Hispanic or Latin origin in the US census while in other countries the term “ethnicity” is a much broader concept encompassing social and cultural characteristics of human populations. Accordingly, applying the HARE approach to international settings or combining cohorts modeled using HARE assignment with others modeled with genetically inferred ancestry groups can create harmonization issues. For instance, meta-analyzing MVP HARE Asian (including Central/South and East Asian in the same population) with Biobank Japan (i.e., East Asian individuals) can lead to a reduction of statistical power. Similarly, applying the HARE assignment to the UK Biobank cohort will lead to a different classification than the ones obtained when applying HARE to the MVP cohort, because of the differences in the SIRE classification in UK and US. The 1kGP+HGDP-based ancestry assignment is based only on genetic information and therefore serves as an international reference panel that can be applied for harmonized classification of cohorts recruited in different parts of the world. Indeed, our 1kGP+HGDP-based ancestry assignment perfectly overlaps with that performed by the Pan-Ancestry analysis recently done in the UK Biobank (see https://pan.ukbb.broadinstitute.org/). The consistent ancestry assignment performed in these cohorts therefore permit valid meta-analysis across two of the largest genetic data repositories in the world in a harmonized fashion. A harmonized definition of ancestry across datasets reduces heterogeneity across the meta-analyzed datasets, increasing the statistical power of the gene discovery analysis.^27; 28^ However, population stratification among the recently-admixed American groups (e.g., African Americans and Latin Americans) still requires careful consideration for within-population adjustment of ancestry diversity.^4–6^

We demonstrated statistically significant improvements in the modeling of ancestry diversity in the MVP cohort, but our study has limitations to consider. First, the MVP is a unique cohort whose diversity reflects many cultural changes through US history filtered through a lens of military service.

Over the 20^th^ century, legislation and military policies gradually expanded opportunities for participation in the US military for women, people of color, and people with different sexual orientations and gender identities. Though demographics of the US military correspond broadly to similar changes in the general population of the US, it remains unclear if they are truly representative, and if our observations reflect directly comparable changes in ancestry proportions across the United States. Second, our findings rely on height as a model phenotype to investigate population stratification biases. Accordingly, the scenarios investigated may differ from those that would be seen for other phenotypes such as medical outcomes that are associated with cultural characteristics and shifts in their recognition, diagnosis, and treatment, possibly introducing additional opportunities for confounding by birth cohort. Third, our study applies a standard set of covariates to adjust for population stratification within each GWAS (e.g., 10 PCs). The inclusion of additional PCs on a case-by-case basis may further reduce evidence of unaccounted-for population stratification, but this may be a costly adjustment resulting in over-correction of test statistics in the GWAS model.

Despite these limitations, our study quantifies the ancestry diversity of MVP HARE superpopulations and demonstrates clear changes in ancestry proportions over time. We further demonstrate the effects of this unaccounted-for ancestry diversity on GWAS and propose one feasible approach to mitigate these effects while still boosting diversity in genetics research. These data are critical for large genetic and meta-analytic studies of health and disease.

## Acknowledgements

This research is based on data from the Million Veteran Program, Office of Research and Development, Veterans Health Administration, and was supported by award # MVP029. This publication does not represent the views of the Department of Veteran Affairs or the United States Government.

## Supplementary Figures

**Figure S1.**
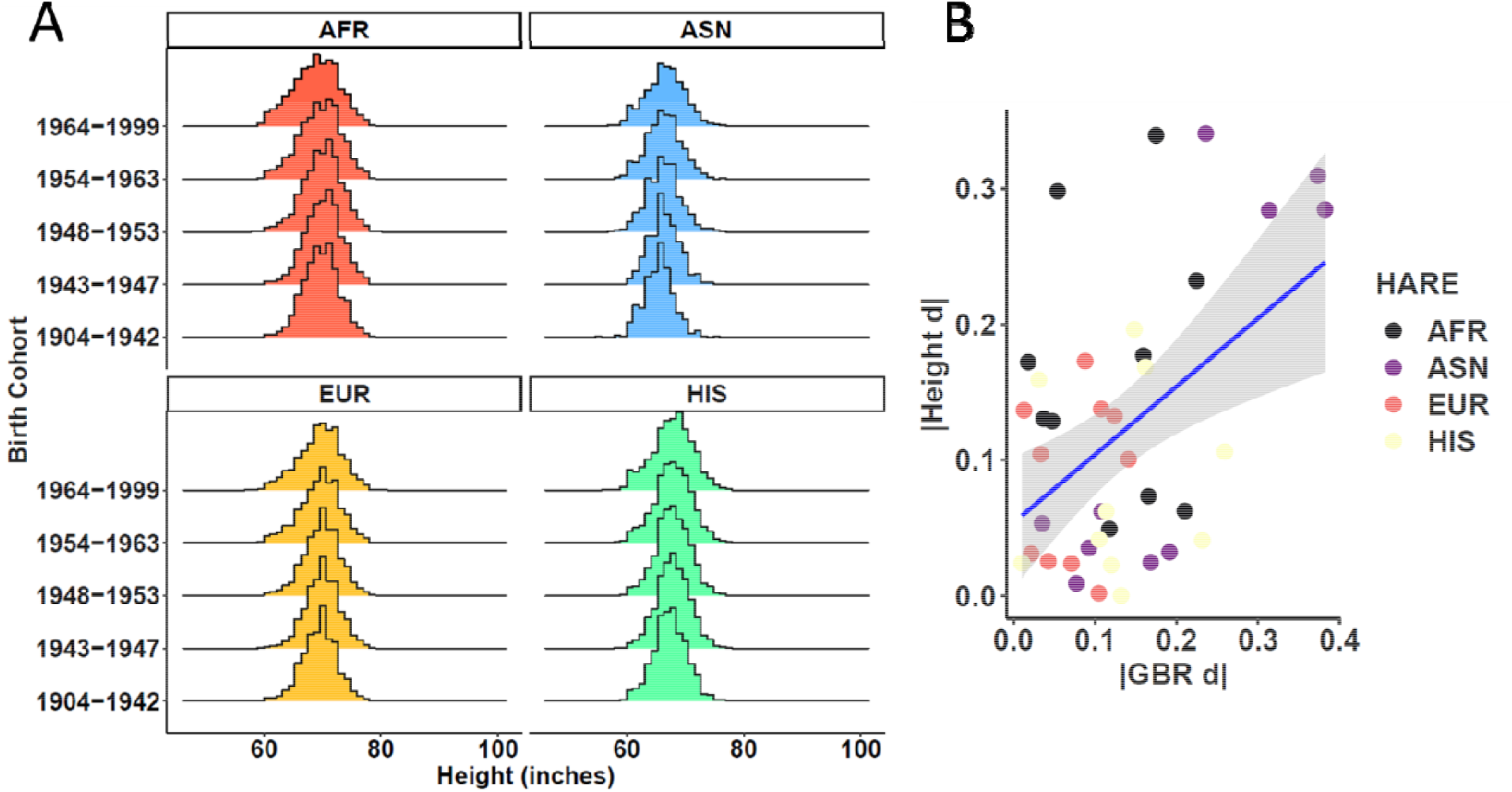
Height across birth cohorts. (A) small changes in height exist across birth cohorts in the MVP. (B) The change in height between two birth cohorts correlates with change in mean GBR ancestry proportion. Each data point is a pairwise comparison of GBR ancestry proportion within each HARE superpopulation. Abbreviations: European (EUR); African (AFR); East Asian (EAS); Hispanic (HIS).

**Figure S2.**
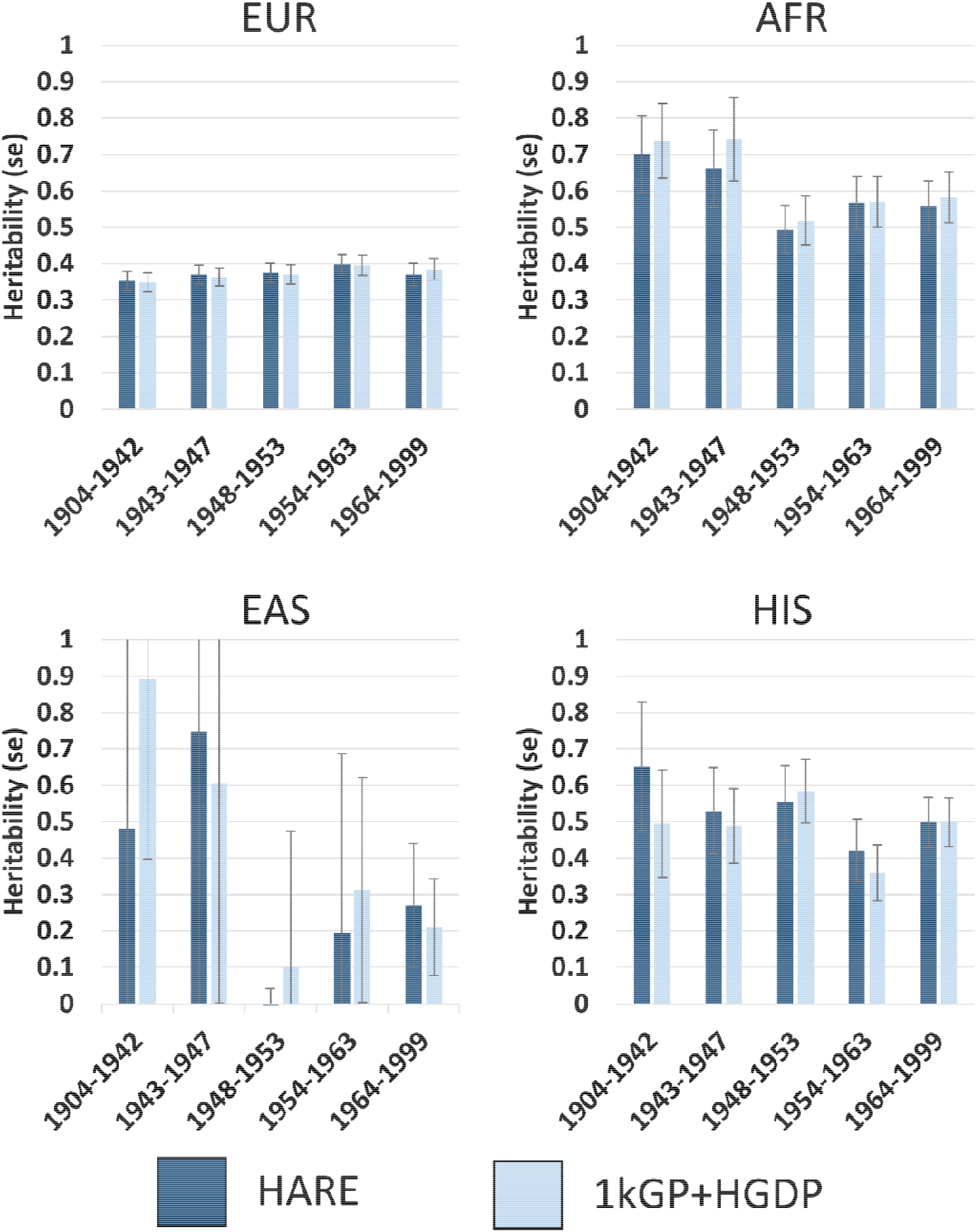
SNP-heritability (h^2^) comparisons across GWAS of height performed in HARE superpopulations and populations assigned using a high-resolution ancestry reference panel composed of 1000 Genomes Project plus Human Genome Diversity Project individuals (1kGP+HGDP). Each GWAS was performed in unrelated participants of the indicated ancestry with age, sex, and 10 withing-population principal components as covariates. Abbreviations: European (EUR); African (AFR); East Asian (EAS); Hispanic (HIS).

**Figure S3.**
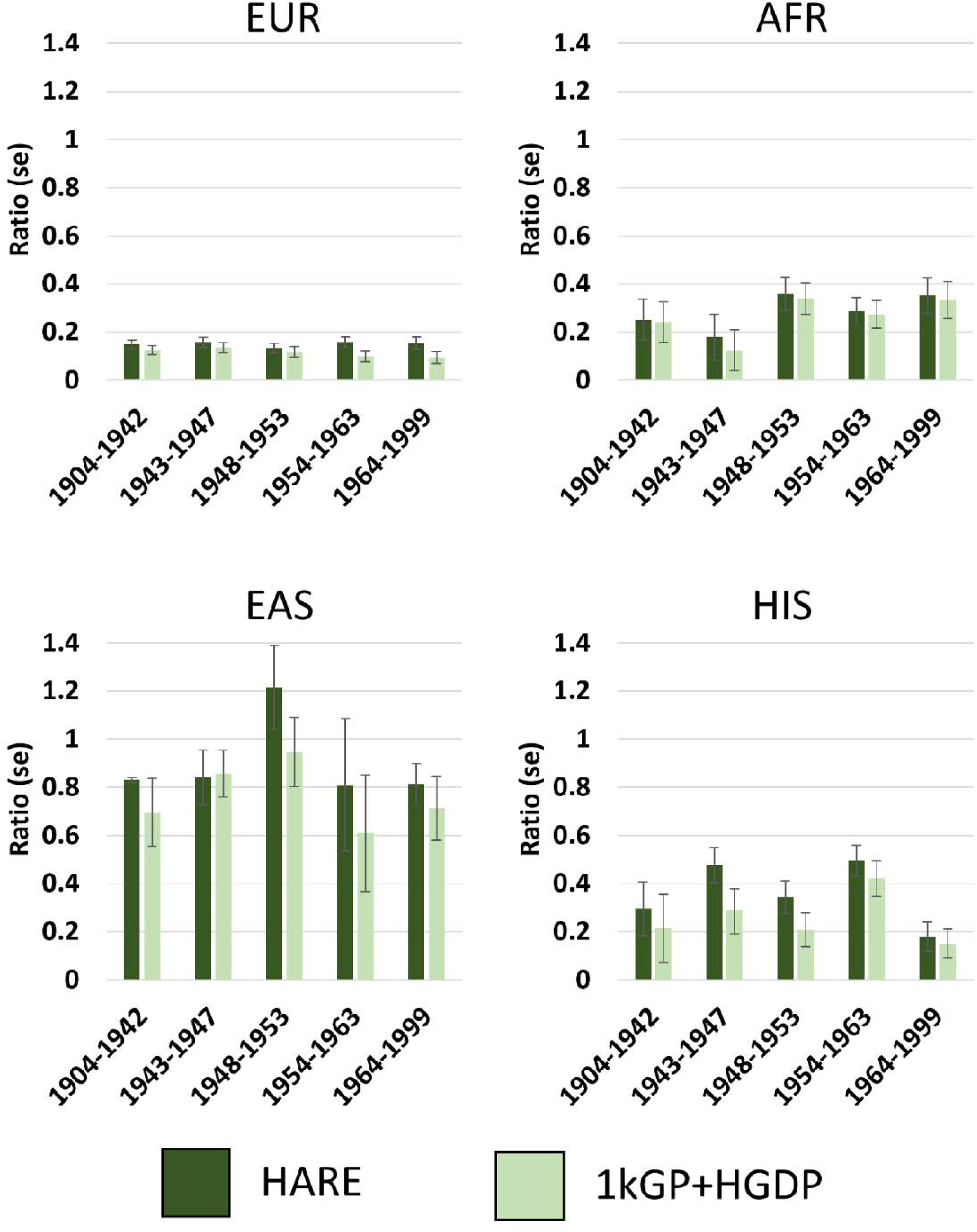
Attenuation ratio comparisons across GWAS of height performed in HARE superpopulations and populations assigned using a high-resolution ancestry reference panel composed of 1000 Genomes Project plus Human Genome Diversity Project individuals (1kGP+HGDP). Each GWAS was performed in unrelated participants of the indicated ancestry with age, sex, and 10 withing-population principal components as covariates. Abbreviations: European (EUR); African (AFR); East Asian (EAS); Hispanic (HIS).

## Supplementary Tables

Table S1. Patterns of service era per birth cohort (“Era”) and across all MVP participants stratified by sex and HARE superpopulations. Each row represents a distinct pattern of service across nine service eras; the frequency of each is calculated by birth cohort and for all MVP participants. Service patterns with less than 11 participants were omitted to preserve data privacy of the participant so HARE total population sample sizes are slightly lower than those reported in Table 1.

Table S2. Sample size per birth cohort derived from cumulative distribution function of year of birth.

Table S3. Mean ancestry proportion of five 1kGP reference populations in all birth cohorts and HARE superpopulations. Two-sided Z-tests were used to compare the statistical difference in means between groups and the corresponding p-values reflect this difference. Standardized mean differences (Cohen’s d) reflects the magnitude of effect size difference between two groups.

Table S4. Comparison of height across birth cohorts in each MVP HARE superpopulations.

Table S5. Metrics for GWAS of height in each ancestry per birth cohort using both methods of population assignment. Heritability (h2), LDSC intercepts, and attenuation ratios were compared across birth cohorts, within each method, using two-sided Z-tests. Multiple testing correction was applied using a false discovery rate of 5%; differences surviving multiple testing correction are highlighted in yellow.

Table S6. Metrics for GWAS of height compared across method used to define superpopulations. Two-sided Z-tests were used to compare heritability (h2), LDSC intercepts, and attenuation ratios between HARE and 1kGP+HGDP superpopulation assignments. Multiple testing correction was applied using a false discovery rate of 5%.

